# The pupal stage shapes life-history trade-offs across metamorphosis

**DOI:** 10.64898/2026.11.02.763434

**Authors:** Kanika Rawat, Navina Mable Francis, Sukanya Samaddar, Kavita Isvaran

## Abstract

Dealing with ecological stressors during development often comes at the expense of later-life performance. Although such trade-offs are well studied as two-stage systems such as larva-adult trade-offs, they are rarely examined across multiple developmental stages, limiting our understanding of how ecological effects are integrated throughout complex life cycles. We addressed this gap using a multistage experimental framework in *Aedes aegypti*, where we exposed individuals to predation risk during the larval stage, the pupal stage, both stages, or neither, and quantified developmental, physiological, and adult fitness traits. Contrary to expectations, predation risk across successive stages did not produce cumulative fitness costs, and traits varied in their responses. Both males and females emerged faster when exposed to risk, with lighter bodies, but males paid an additional cost of reduced starvation resistance when the pupal stage was exposed. Surprisingly, despite their lower body mass, females showed enhanced reproductive performance when both stages were exposed to risk. Overall, responses were stage- and sex-specific, with exposure during the non-feeding pupal stage imposing greater life-history costs than larval exposure alone. Measurements across the larva-pupa-adult continuum further revealed that traits such as body mass and energy reserves respond differently before and after metamorphosis, highlighting the role of the pupa in mediating trade-offs. Our findings identify the pupal stage as a critical mediator of life-history trade-offs and demonstrate the value of a multistage framework for understanding the role of each stage in life-history evolution.

## Introduction

Organisms can acquire or process only limited resources, which must be allocated among multiple traits to maximise fitness. Due to shared resources between various traits, increased investment in one trait often comes at the expense of another, a phenomenon known as inter-trait trade-offs (Fabian and Flatt 2012, Garland Jr 2022, Martin and Bize 2018, Reznick 1985, Stearns 1989, Stearns 2000). As organisms transition through their life cycle, trade-offs can occur between early and late life stages due to shared resources across the life cycle. For example, trade-offs between rapid body growth leading to early reproduction vs. increased survival and reproduction later in life have been reported in various vertebrate taxa (reviewed in Lemaître et al. 2015).

Many life cycles observe drastic transitions before adulthood, known as metamorphosis, with distinct life stages characterised by unique morphologies and functions and varied ecology (Wilbur 1980). Ecological pressures that drive increased investment towards a trait in one stage can result in trade-offs affecting subsequent stages in such complex life cycles (Pechenik 2006). For instance, damselfly (*Coenagrion puella*) larvae that invest in cuticular melanin under UV exposure develop into adults with reduced immune response (Debecker et al. 2015). Other examples include reduced development time in common frog (*Rana temporaria*) tadpoles to evade decreasing water levels, trading off against body size at emergence (Johansson et al. 2010), and increased melanisation in wood tiger moth (*Arctia plantaginis*) larvae under predation risk at the cost of reduced pupal body weight (Almeida et al. 2021).

Most trade-off studies in complex life cycles primarily focus on traits within a single stage (e.g., fecundity vs. survival) or between two stages (e.g., growth vs. body size), even though taxa, including cnidarians, insects, crustaceans, echinoderms and molluscs, have multiple distinct life stages. While examining trade-offs between two stages provides valuable insights into how one stage influences the other, it often overlooks the ecological roles of other stages. This gap limits our understanding of how each stage contributes to overall fitness through its share of resource utilisation and ability to manage trade-offs.

Holometabolous insects, among the most diverse and successful lineages (Kristensen 1999, Rolff et al. 2019, Whiting 2004), provide an excellent opportunity to investigate multistage trade-offs. Their life cycles are marked by a transition stage between larva and adult, known as pupa. Existing trade-off studies primarily emphasise how larval ecology influences adult traits, as larval conditions significantly impact fitness due to their roles in nutrition and growth (Wilbur, 1980). The larval stage is a voracious feeding phase crucial for acquiring resources necessary for growth during metamorphosis and supporting adult maintenance and reproduction (Aguila et al. 2013, Boggs and Freeman 2005, Colasurdo et al. 2009, Poças et al. 2022). Therefore, various ecological factors that interfere with larval nutrition and function affect individual fitness. For example, restricted resource availability during the larval stage may reduce investment in spermatogenesis and decrease sperm production in meal moths, *Plodia interpunctella* (Gage and Cook 1994). Similarly, high larval density reduces adult longevity and fecundity in the Mediterranean fruit fly, *Ceratitis capitata* (Diamantidis et al. 2020). Predation stress during the *Anopheles gambiae* larval stage can alter nutrient allocation, reduce adult body size, and cause oxidative damage to adult biomolecules (Roux et al. 2021). Heat stress during the larval stage reduces feeding and lowers lipid reserves in *Drosophila melanogaster* adults (Karpova et al. 2024).

The pupal stage has been largely overlooked in trade-off studies. This neglect may stem from it being a non-feeding stage. However, this trait makes pupa a crucial and sensitive link in the life cycle, as it must transition into an adult using a fixed amount of energy while managing trade-offs under persistent ecological stressors. For example, heat stress during the pupal stage has been shown to affect life-history traits—including fecundity, body size and longevity—in various insect species such as ladybirds, hoverflies, parasitoid wasps, and the oriental fruit fly (Awde et al. 2023, Daňková et al. 2023, El-Sabrout et al. 2021, Mutamiswa et al. 2023). Additionally, despite being traditionally viewed as a quiescent stage, pupae in many insect species exhibit behavioural defences such as vibrations and abdominal rotations (Hinton 1946, Ichikawa and Sakamoto 2013, Lindstedt et al. 2019, Singh et al. 2023). These behaviours are effective against various threats (Eisner and Eisner 1992, Kojima et al. 2012). For instance, abdominal rotations by pupae of tenebrionid beetles are effective against cannibalistic larvae (Ichikawa and Kurauchi 2009). As pupae cope with ecological stressors, their physiological and behavioural responses may involve trade-offs with consequences for later stages. This makes the pupal stage essential for investigating life-history trade-offs and understanding how early ecological conditions shape trait expression across the larval-pupal-adult continuum.

To explore how trade-off consequences accumulate across multiple stages and the role of each stage in managing trade-offs, we studied the effects of predation risk and its fitness consequences across three stages of the mosquito *Aedes aegypti*. Multiple studies on mosquito species have shown that predation risk during the larval stage imposes costs on adult survival, body size and fecundity (Beketov and Liess 2007, Chandrasegaran and Juliano 2019, Ower and Juliano 2019). Such costs likely arise from larval behavioural responses, such as reducing activity/conspicuousness compromising nutritional intake (Ball and Baker 1996, Costanzo et al. 2011, Lima and Dill 1990, van Uitregt et al. 2012), or physiological stress responses (Adamo and Baker 2011, Janssens and Stoks 2013, Roux et al. 2021, Slos and Stoks 2008), including early emergence in some cases (Alcalay et al. 2018, Ower and Juliano 2019). Therefore, adult traits are negatively affected, particularly under low food availability (Chandrasegaran et al. 2018).

In addition, a few studies on mosquito pupae have revealed that they respond to disturbances and predators in their microhabitats by modifying their space use and diving behaviour (Awasthi et al. 2012, Chandrasegaran et al. 2020, Romoser 1975, Romoser and Lucas 1999). Their diving response is also known to incur energy costs and reduce adult survival (Lucas and Romoser 2001). These pupal responses to predation risk could trade off against adult traits, but such relationships have rarely been explored previously. Thus, *Aedes aegypti* provides an ideal system for investigating how ecological stressors, specifically predation risk, experienced during both larval and pupal stages translate across life stages to affect overall fitness.

We studied the independent and combined effects of larval and pupal predation risk exposure on key life-history traits, including development time, size, survival, and reproductive success. Since *Aedes aegypti* larvae and pupae share the same habitat and similar threats, we examined trade-offs along the larva-pupa-adult continuum driven by larval and pupal responses to predation risk. We hypothesised that trade-off consequences would be more pronounced when both larval and pupal stages are exposed to predation risk than when only one stage is exposed. This is because each stage is expected to allocate resources towards responding to predation threats, resulting in a cumulative cost for the adult stage. Accordingly, we predicted that key life-history traits such as survival would be lowest when both stages face predation risk, intermediate when one stage is exposed, and highest when neither stage experiences risk.

To test this hypothesis in a comparative framework, we established four larval-pupal environmental combinations: (1) no risk exposure during either stage (L-P-), (2) larva exposed to risk (L+P-), (3) pupa exposed to risk (L-P+), and (4) both stages exposed to risk (L+P+). Across these groups, we measured and compared a comprehensive set of life-history traits: survival to adulthood, time to emergence, adult wet weight or mass, resistance to starvation, female reproductive success, and oviposition time. Additionally, we estimated energy reserves to assess physiological-level trade-offs and infer life-history outcomes that could not be directly measured. We also examined how the larval environment influences pupal traits to isolate risk effects up to the pupal stage and gain insights into the pupa’s role in mediating trade-offs that ultimately affect adult traits.

## Methods

### Predator collection and mosquito rearing

We collected *Bradinopyga geminata* (granite ghost dragonfly) nymphs, predators of *Aedes aegypti* larvae and pupae, from rock pools around Bengaluru (2022-2023). The nymphs were stored in small containers filled with tap water and fed four to six mosquito larvae or pupae. To maintain water quality, we replaced the water in the containers every other day. We reared *Aedes aegypti* mosquitoes in the lab colony under a 12:12 hours light:dark cycle, with 60-75% relative humidity and 24-27 °C temperature. Adult males and females were housed in 30 x 30 x 30 cm cages where they were allowed to mate. Females were fed human blood using a blood-feeding system, while both sexes were provided with diluted honey on cotton pads as a sugar source. The females laid eggs in plastic cups with their walls lined with blotting paper. We dried these eggs, stored them, and later hatched one-to three-month-old eggs in enamel trays filled with tap water for the experiment.

### Experimental setup

Twenty-four hours post-hatching, we counted and distributed 45 first instar larvae into each tray containing two litres of tap water with 0.15 g of larval food. The food level and larval density were selected based on preliminary experiments assessing larval development across a range of food conditions. This combination was chosen to reflect resource competition in the rearing environment, as mosquito larvae typically develop in nutrient-limited, competitive habitats in the wild (Arrivillaga and Barrera 2004, Barrera et al. 2006). We established four predation-risk treatments to examine the effects of stage-specific exposure:

- **L-P-**: Neither larval nor pupal stage exposed to predation risk
- **L+P-**: Larval stage exposed to predation risk, pupal stage not
- **L-P+**: Larval stage not exposed to predation risk, pupal stage exposed
- **L+P+**: Both larval and pupal stages exposed to predation risk

We placed two predator nymphs (one per 50 mL transparent container) in each predation-risk tray, while the no-risk trays contained two empty containers each. The nymphs were fed daily with six larvae (third and fourth instars) and pupae from the mosquito colony. To expose the growing larvae to predation cues, i.e., dead remains of larvae and pupae and predator kairomones, we mixed the water in the predator containers and trays twice daily. We also mixed water in the no-risk trays to maintain consistent handling for all treatments. As larvae developed and grew big, we used perforated predator containers to ensure continued and effective passage of predation cues through the pores of the containers.

Around the sixth day of the larval phase, we started looking for larvae metamorphosing into pupae at six-hour intervals. As soon as a few pupae were spotted in a tray, we rinsed the remaining larvae twice with water to remove residual predation cues and transferred them to another tray based on the designated treatment combination. For example, larvae from L+P- and L-P-trays were moved to no-risk trays to ensure that the soon-metamorphosing larvae were not exposed to risk as pupae. In contrast, larvae from L+P+ and L-P+ trays were transferred to trays containing predation cues.

The experimental conditions ranged from 24-27 °C in temperature and 50-75% in humidity across all trials.

### Measurement of traits from pupal to adult stage (Fig 1a)

#### Pupal survival to adulthood and time to emergence

After approximately 48 hours of pupation, we transferred pupae to 1.5 mL vials and monitored adult emergence at six-hour intervals. Individuals that emerged as adults were recorded for their sex and time of emergence. Pupal survival was calculated as the proportion of pupae surviving to adulthood by dividing the number of individuals that emerged by the total number of pupae in vials. Emergence time (from hatching) was also measured for each pupa. We then randomly selected subsets of adults to measure starvation resistance, wet weight and energy reserves, and reproductive success and oviposition time (in females). Sample sizes per treatment and trait are provided in Table 1.

**Fig 1:**
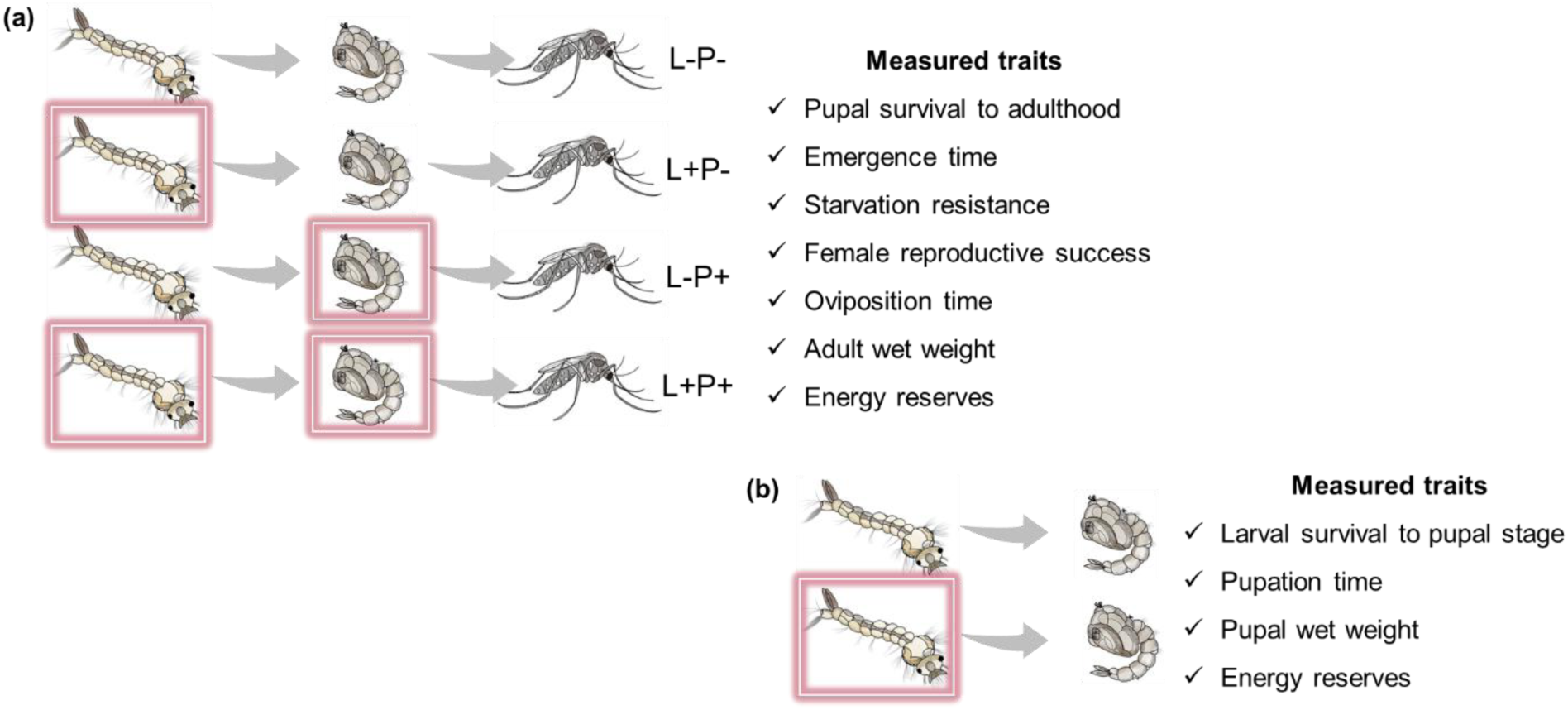
Experimental design: (a) Larvae and pupae were reared under different combinations of predation-risk environments, resulting in four treatments: no risk exposure at either stage (L-P-), risk exposure during the larval stage (L+P-), risk exposure during the pupal stage (L-P+), and risk exposure during both stages (L+P+). We measured traits during the pupal-to-adult transition and in the adult stage to assess both stage-specific and cumulative effects of predation risk. (b) To isolate the effects of larval-stage risk up to the pupal stage, we compared traits during the larva-to-pupa transition and in the pupal stage across risk and no-risk environments. Red boxes indicate the presence of predation risk.

**Table 1:** Treatment-specific sample sizes for analyses of adult traits. F = female; M = male.

| **Trait** | **Sample size per treatment** |  |  |  |  |  |  |  |
|---|---|---|---|---|---|---|---|---|
|  | L-P- |  | L+P- |  | L-P+ |  | L+P+ |  |
|  | F | M | F | M | F | M | F | M |
| Emergence | 202 | 172 | 157 | 110 | 188 | 155 | 241 | 173 |
| Wet weight | 123 | 123 | 99 | 105 | 153 | 153 | 135 | 129 |
| Starvation resistance | 107 | 97 | 100 | 81 | 90 | 45 | 149 | 98 |
| Protein reserve | 36 | 37 | 28 | 30 | 45 | 45 | 39 | 37 |
| Glycogen reserve | 35 | 34 | 27 | 29 | 45 | 45 | 39 | 37 |
| Lipid reserve | 36 | 37 | 28 | 30 | 45 | 44 | 38 | 37 |

#### Starvation resistance

We maintained adults on water only, without food, in perforated 1.5 mL tubes to ensure adequate ventilation and monitored them every 12 hours to record their time of death. Total hours from hatching until death were used as an index of starvation resistance or lifespan of starved individuals.

#### Reproductive success and oviposition time

We tracked oviposition for 24-26 days following the emergence of females from different predation-risk treatments. This period represents most of the adult lifespan since *Aedes* females typically live 25-50 days (median survival time) under optimal laboratory conditions (Goindin et al. 2015, Posidonio et al. 2021, Villarreal et al. 2018, Yeap et al. 2011), with fecundity declining as they age (Maciel-de-Freitas et al. 2011, Ramírez-Sánchez et al. 2023).

We introduced eight females per treatment in a 15 x 15 x 15 cm cage after 1-2 days of their emergence, along with four untreated males from the colony. They were allowed to mate and provided with a 10% sucrose solution (10 mL soaked in a cotton pad) from the following day onward, which was replenished every three days. Females underwent eight blood-feeding cycles, with successive blood-feeding episodes separated by two days. We provided blood meals using a membrane-feeding system in which blood was held in glass feeders maintained at human body temperature by a surrounding water jacket. The feeders were sealed with a Parafilm membrane, through which females fed when the feeders were placed on top of the cages. Males were removed from the cages after a week, i.e., two blood-feeding cycles. We provided 2 mL of blood per cage from 11 AM to 3 PM and kept a 50 mL container lined with water-soaked blotting paper as an oviposition substrate, which was replaced every blood-feeding cycle. Eggs laid on the blotting paper (ovistrip) were counted, and any female mortality in the cages was recorded every cycle.

The proportion of surviving females and the number of eggs per surviving female per cycle were used to calculate lifetime reproductive success and oviposition time. Oviposition time for an egg was measured as the time from the day cages were set up (1-2 days after female emergence) until the egg was deposited on the ovistrip. Female reproductive success was calculated as an average for each cage of eight females since eggs from a cage could not be assigned to individual females.

#### Adult wet weight and energy reserves

We stored the adults at −20 °C for their energy reserves estimation. Before performing the biochemical assays to estimate protein, glycogen and lipid, individuals were thawed on ice, blotted dry with tissue, and weighed using a microbalance with a readability of 0.01 mg. We then added 60 µL of 2% sodium sulphate solution to each tube containing one individual and homogenised each sample using a mechanical bead beater (Qiagen-TissueLyser II) at a frequency of 25 Hz (three cycles of 20 seconds with 5-second pauses). 50 µL of homogenate was divided into 7.5 µL and 42.5 µL portions for protein and glycogen-lipid quantification, respectively, adapting the protocols mentioned in Chandrasegaran et al. 2018, Foray et al. 2012, van Handel 1985a, van Handel 1985b.

For protein estimation, we diluted 7.5 µL of the sample homogenate with 42.5 µL of 2% sodium sulphate. 400 µL BCA reagent was added to the diluted sample following the Millipore (71285-3 BCA protein assay) kit’s protocol and incubated for two hours. We measured the absorbance of the well-mixed final product at 562 nm using BSA in 2% sodium sulphate as standard, prepared using the same kit.

The 42.5 µL of the sample was mixed with chloroform-methanol (1:2 v/v) solution to a final volume of 425 µL. We aliquoted 275 µL of that for glycogen estimation and 150 µL for lipid estimation. For glycogen estimation, we centrifuged the 275 µL sample-chloroform-methanol mixture (4 °C, 13,000 rpm, 2 minutes), discarded the supernatant, air-dried the pellet, and added 1 mL of anthrone to it. We vortexed the tube contents to dissolve the pellet and heated the mixture at 90 °C in a water bath for 90 seconds. The mixture was immediately kept on ice, mixed well, and absorbance was measured at 625 nm using glucose in distilled water as standard.

For lipid estimation, we evaporated the 150 µL sample-chloroform-methanol mixture in a dry bath (5-10 minutes), cooled it, and mixed it with 20 µL sulphuric acid. The mixture was heated for 2 minutes in a 90 °C water bath, kept on ice, and mixed with 480 µL vanillin. We incubated the mixture for 10 minutes and measured absorbance at 525 nm using vegetable oil in chloroform as standard. The estimated energy reserves were normalised to the wet weight of each individual.

### Measurement of traits from larval to pupal stage (Fig 1b)

#### Larval survival to pupal stage and time to pupation

We inspected trays at six-hour intervals for pupation, starting on the sixth day of the larval phase. To track the average time to pupation in trays with and without larval-stage risk exposure, we meticulously recorded the number of pupae per tray at each interval, up to two days after the first pupation event (i.e., until at least 95% of larvae pupate). We divided the total number of pupae and the remaining surviving larvae per tray by 45 to calculate the number of larvae that survived until 95% of larvae had pupated. We also used the pupation time of individual pupae to compare the average pupation time between risk and no-risk environments during the larval stage.

#### Pupal wet weight and energy reserves

While one set of trays was dedicated to recording pupation, another was used to collect individuals at pupation for freezing and subsequent measurement of wet weight and energy reserves, following the same protocol described above for adults. The sample size per treatment for each trait is provided in the respective figure legends (Fig 6 and Fig 7).

### Statistical analyses

Statistical analyses were conducted in R version 4.2.3. We used appropriate linear, generalised linear, and mixed-effects models to test for differences among predation-risk treatments while avoiding multiple pairwise t-tests. Standard treatment contrasts in linear models typically compare each level to a single reference group, which limits the ability to customise comparisons of interest. To address this, we employed orthogonal contrasts, which allow for the specification of statistically independent and interpretable comparisons (Crawley 2012). For the four-level treatment design (L-P-, L+P-, L-P+, L+P+), we defined three orthogonal (statistically independent) contrasts. The first contrast tested the overall effect of predation risk by comparing the no-risk treatment (L-P-) to the average of all risk-exposed treatments (L+P-, L-P+, L+P+). The second contrast compared the average risk effect of single-stage exposure (L+P-and L-P+) to both-stage exposure (L+P+), assessing whether risk experienced across both developmental stages had a stronger effect than risk at one stage. The third contrast tested for differences between the risk effects of larval-only and pupal-only risk exposure (L+P-vs. L-P+), thereby isolating the relative contributions of each developmental stage to trait variation.

Separate models were set up for male and female traits. We included experimental blocks as fixed effects due to the small number of blocks, which made it inappropriate to treat them as random effects (Bolker et al. 2009). For traits averaged over trays or cages, such as reproductive traits, blocks were kept as random effects due to limited sample sizes. The results from these models were qualitatively similar to those where blocks were included as fixed effects.

Specific models were selected according to the nature of each trait or response variable to analyse differences across the orthogonal contrasts. Pupal survival to adulthood, best represented as a binomial variable, was analysed using a generalised binomial model with a logit link. Time to emergence, wet weight, and starvation resistance were compared using linear models. Reproductive success and oviposition time were analysed using linear mixed-effects models. Energy reserves were compared using linear mixed-effects models, incorporating assay day as a random effect to account for variability arising from manual handling.

Similarly, we compared traits measured during the larva-to-pupa transition between larval-risk and no-risk environments. Time to pupation was analysed using a linear model, while larval survival to the pupal stage (recorded as alive/dead) was evaluated using a generalised mixed-effects binomial model. Energy reserves were analysed using linear mixed-effects models, with assay day included as a random effect.

As an additional check for the robustness of the results, all models were evaluated using confidence intervals generated through bootstrap routines. The bootstrapping analyses largely supported the results from the statistical models. We visualised the data as means with 95% bootstrapped confidence intervals based on the original dataset, offering insight into effect sizes. Block effects were accounted for by subtracting them during the bootstrap process to improve visualisation.

Inferences were primarily based on statistically detectable effects indicated by model results, supported by patterns in means and 95% bootstrapped confidence intervals from the original data, which reflect both the magnitude of estimates and the uncertainty around them. In cases where effects were weakly detectable—i.e., when the bootstrapped confidence interval for one treatment did not overlap the mean of another, but the model-derived confidence interval for their direct comparison slightly overlapped zero—we included a description of these findings. However, such marginal or weak effects were not emphasised or discussed further.

## Results

### Effects of larval-pupal environments on traits

#### Pupal survival to adulthood and time to emergence

Pupal survival or the proportion of pupae that survive to adulthood was similar across predation-risk treatments (Fig 2, Table 2). However, the average emergence time varied across risk treatments. On average, predation risk during the immature stages accelerated emergence for females and males by 3-4 hours (Fig 3a, Table 3: Time to emergence, contrast 1). We did not detect any difference in time to emergence among the stage-specific risk environments in males (Table 3: Time to emergence, contrasts 2 & 3). In females, the effect of exposure to predation risk during both larval and pupal stages did not differ statistically from the pooled effects of single-stage risk exposure (Table 3: Time to emergence, contrast 2). However, within single-stage risk environments, emergence time varied depending on the stage of risk exposure. Females exposed to predation risk during the larval stage emerged 1-2 hours earlier than those exposed during the pupal stage (Fig 3a, Table 3: Time to emergence, contrast 3). Therefore, although the pooled single-stage vs. both-stage risk exposures (contrast 2) showed similar mean effects, larval-stage risk exposure—unlike pupal-stage exposure—had a different effect from both-stage exposure. As suggested by means and bootstrapped confidence intervals, risk exposure during both stages (L+P+) slightly delayed emergence (by 1.5 hours) compared with risk exposure exclusively during the larval stage (L+P-) (Fig 3a).

**Fig 2:**
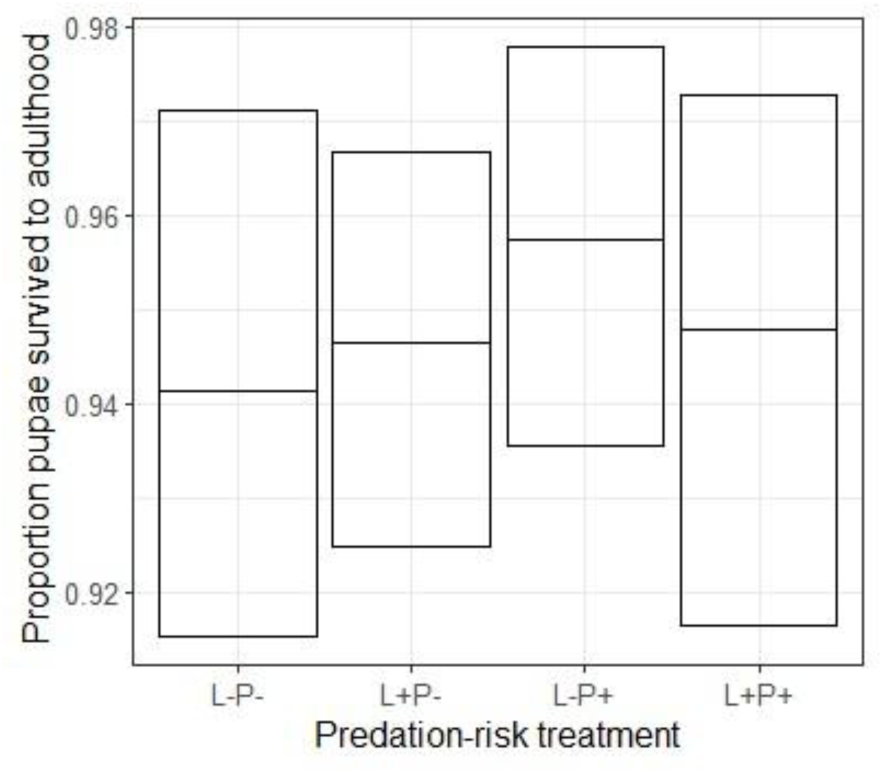
Mean proportions of pupae surviving to adulthood with 95% bootstrap confidence intervals across different risk treatments. Sample sizes for each treatment: no-risk (L-P-, n = 377, 10 trays), larval-stage risk (L+P-, n = 239, 6 trays), pupal-stage risk (L-P+, n = 343, 9 trays), and both-stage risk (L+P+, n = 379, 10 trays).

**Fig 3:**
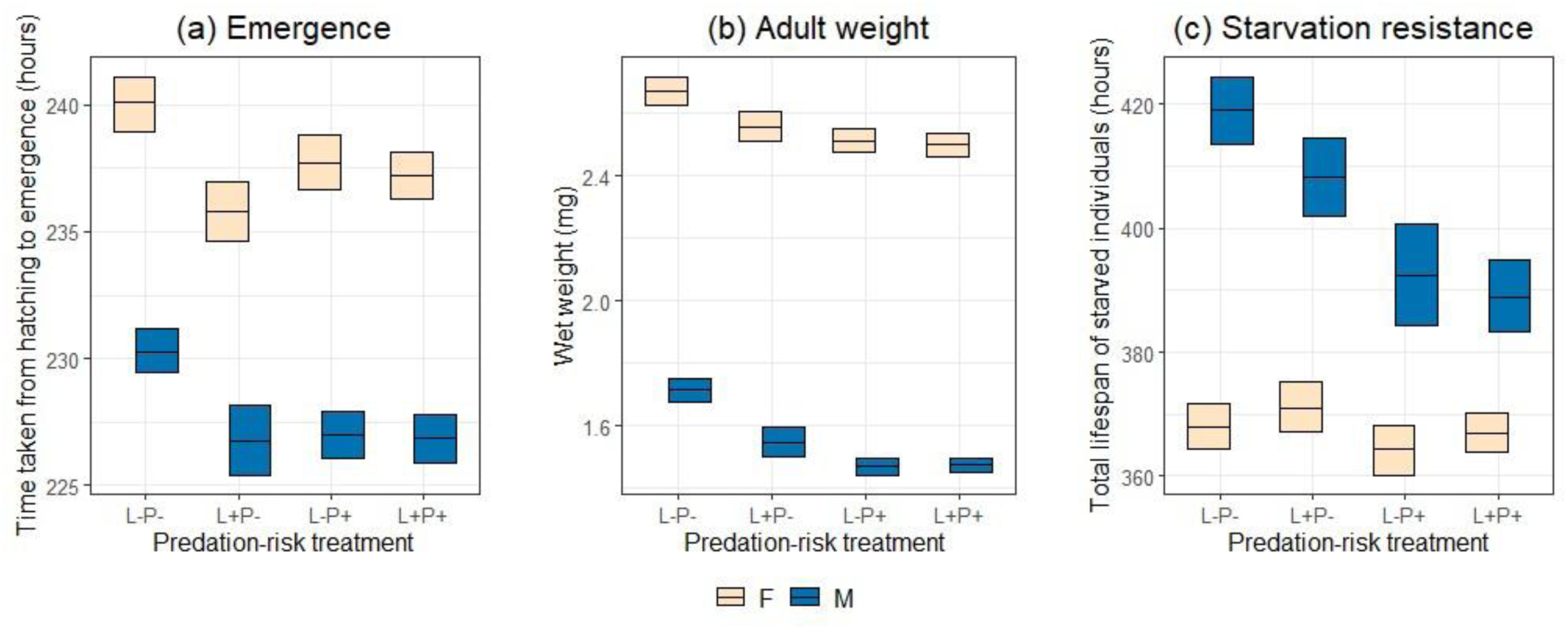
(a) Emergence, (b) weight and (c) starvation resistance data (mean + 95% bootstrap confidence intervals) for females (ivory) and males (blue) from different risk treatments, i.e., no-risk (L-P-), risk during the larval stage (L+P-), the pupal stage (L-P+), and both the stages (L+P+). Sample sizes: emergence (F = 788, M = 610), weight (F = 510, M = 510), and starvation resistance (F = 446, M = 321). Treatment-specific sample sizes are provided in Table 1. The presentation facilitates direct comparison of means using 95% bootstrap confidence intervals from the data.

**Table 2:**
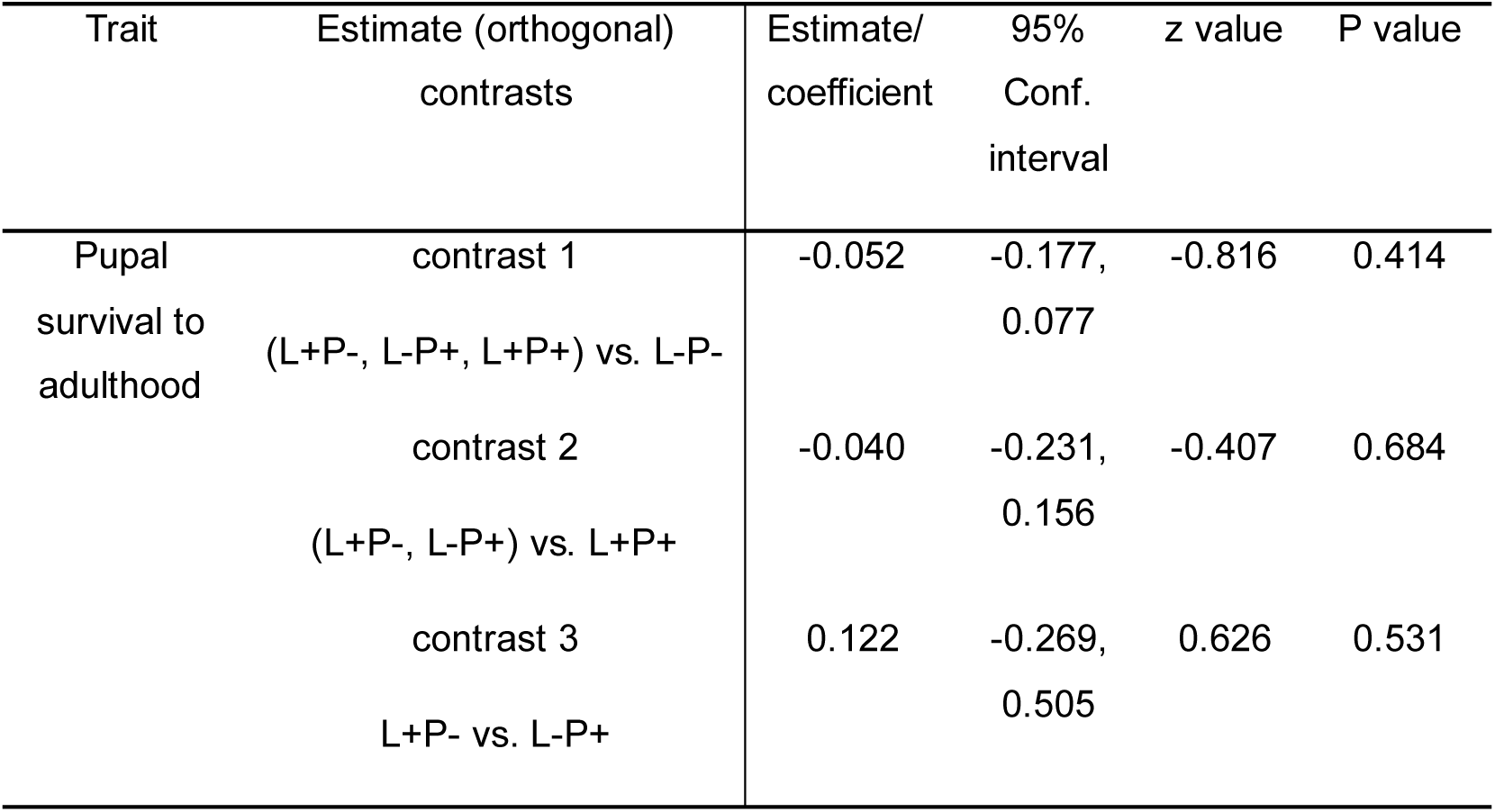
Model estimates, their confidence intervals, and test statistics from the binomial generalised linear model, showing differences in the proportion of pupae survived to adulthood across different risk treatments, i.e., risk vs. no-risk (contrast 1), risk during one stage vs. both the stages (contrast 2), risk during larval vs. pupal stage (contrast 3).

| Trait | Estimate (orthogonal) contrasts | Estimate/ coefficient | 95% Conf. interval | z value | P value |
|---|---|---|---|---|---|
| Pupal survival to adulthood | contrast 1<br>(L+P-, L-P+, L+P+) vs. L-P- | -0.052 | -0.177, 0.077 | -0.816 | 0.414 |
|  | contrast 2<br>(L+P-, L-P+) vs. L+P+ | -0.040 | -0.231, 0.156 | -0.407 | 0.684 |
|  | contrast 3<br>L+P- vs. L-P+ | 0.122 | -0.269, 0.505 | 0.626 | 0.531 |

**Table 3:**
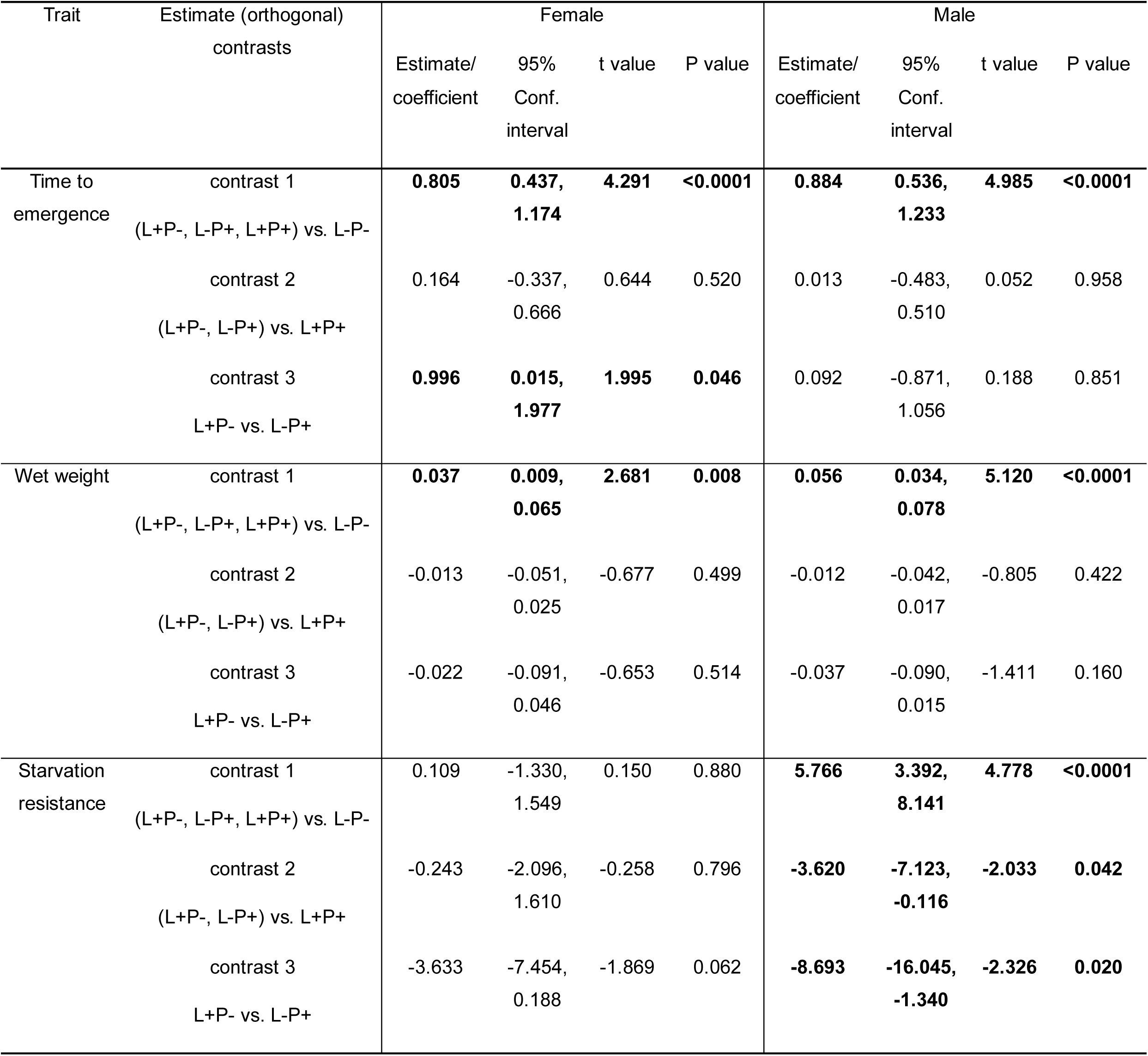
Model estimates, their confidence intervals, and test statistics from separate linear models for females and males, highlighting differences in time to emergence, wet weight, and total lifespan of starved individuals under different risk treatments, i.e., risk vs. no-risk (contrast 1), risk during one stage vs. both the stages (contrast 2), risk during larval vs. pupal stage (contrast 3). Entries in bold denote statistically detectable effects. Differences between males and females can be qualitatively inferred by assessing overlap between 95% confidence intervals and corresponding contrast estimates.

| Trait | Estimate (orthogonal) contrasts | Female |  |  |  | Male |  |  |  |
|---|---|---|---|---|---|---|---|---|---|
|  |  | Estimate/ coefficient | 95% Conf. interval | t value | P value | Estimate/ coefficient | 95% Conf. interval | t value | P value |
| Time to emergence | contrast 1<br>(L+P-, L-P+, L+P+) vs. L-P- | **0.805** | **0.437, 1.174** | **4.291** | **<0.0001** | **0.884** | **0.536, 1.233** | **4.985** | **<0.0001** |
|  | contrast 2<br>(L+P-, L-P+) vs. L+P+ | 0.164 | -0.337, 0.666 | 0.644 | 0.520 | 0.013 | -0.483, 0.510 | 0.052 | 0.958 |
|  | contrast 3<br>L+P- vs. L-P+ | **0.996** | **0.015, 1.977** | **1.995** | **0.046** | 0.092 | -0.871, 1.056 | 0.188 | 0.851 |
| Wet weight | contrast 1<br>(L+P-, L-P+, L+P+) vs. L-P- | **0.037** | **0.009, 0.065** | **2.681** | **0.008** | **0.056** | **0.034, 0.078** | **5.120** | **<0.0001** |
|  | contrast 2<br>(L+P-, L-P+) vs. L+P+ | -0.013 | -0.051, 0.025 | -0.677 | 0.499 | -0.012 | -0.042, 0.017 | -0.805 | 0.422 |
|  | contrast 3<br>L+P- vs. L-P+ | -0.022 | -0.091, 0.046 | -0.653 | 0.514 | -0.037 | -0.090, 0.015 | -1.411 | 0.160 |
| Starvation resistance | contrast 1<br>(L+P-, L-P+, L+P+) vs. L-P- | 0.109 | -1.330, 1.549 | 0.150 | 0.880 | **5.766** | **3.392, 8.141** | **4.778** | **<0.0001** |
|  | contrast 2<br>(L+P-, L-P+) vs. L+P+ | -0.243 | -2.096, 1.610 | -0.258 | 0.796 | **-3.620** | **-7.123, -0.116** | **-2.033** | **0.042** |
|  | contrast 3<br>L+P- vs. L-P+ | -3.633 | -7.454, 0.188 | -1.869 | 0.062 | **-8.693** | **-16.045, -1.340** | **-2.326** | **0.020** |

#### Adult weight and resistance to starvation

On average, females from risk environments weighed 0.1-0.15 mg less, i.e., 4-6% less, and males weighed 0.2-0.25 mg less, i.e., 10-14% less than individuals from the no-risk environment (Fig 3b, Table 3: Wet weight, contrast 1). Furthermore, no consistent differences in mass were detected among the stage-specific risk environments (Table 3: Wet weight, contrasts 2 & 3). A comparison of means and 95% bootstrapped confidence intervals suggested that males exposed to risk during the pupal stage may have emerged even lighter than those from the larval-risk environment. However, this difference was not detected by contrast 3 (L+P-vs. L-P+; *t* = −1.411, *P* = 0.160).

Risk exposure during the immature stages did not substantially affect female resistance to starvation, and no stage-specific effects were detected within the risk environments (Table 3: Starvation resistance, contrasts 1-3). However, a comparison of means and 95% bootstrapped confidence intervals suggested that females exposed to risk during the pupal stage may have survived for a shorter duration than those from the larval-risk environment (Fig 3c), although this difference was only weakly detected by contrast 3 (L+P-vs. L-P+; *t* = −1.869, *P* = 0.062).

In contrast to females, male lifespan under starvation was markedly reduced by 10-30 hours (i.e., by 2-7% relative to L-P-) when exposed to risk during the immature stages (Fig 3c, Table 3: Starvation resistance, contrast 1). The influence of risk on the lifespan of starved males varied with the stage of exposure: exposure during both larval and pupal stages resulted in a more substantial negative effect than the pooled single-stage exposures (Table 3: Starvation resistance, contrast 2). However, a pronounced difference in starvation resistance between the single-stage risk environments (Fig 3c, Table 3: Starvation resistance, contrast 3) indicated that larval-stage risk exposure—but not pupal-stage exposure—drove the difference between the pooled single-stage and both-stage risk effects (contrast 2). Males exposed to risk only during the larval stage survived starvation considerably better, with total lifespans 15-16 hours longer than those exposed during the pupal stage, with or without the larval stage (Fig 3c).

In summary, in males, risk exposure exclusively during the larval stage reduced lifespan under starvation by 10-11 hours, while exposure during the pupal stage resulted in a more pronounced reduction of 25-27 hours—surprisingly, as severe as that caused by both-stage exposure.

#### Reproductive success and oviposition time

On average, early-life risk environments did not produce a detectable effect on the mean reproductive success compared to the no-risk environment (Table 4: Reproductive success, contrast 1). Furthermore, the stage-specific risk environments did not statistically differ in their effects on reproductive success (Table 4: Reproductive success, contrasts 2 & 3). However, comparisons of means and 95% bootstrapped confidence intervals revealed that females exposed to risk during both larval and pupal stages laid approximately 50% more eggs than females from the no-risk environment (Fig 4a). This difference, however, could not be detected statistically through contrast 1 because the reproductive success of females from the single-stage risk environments (L+P-and L-P+) was statistically indistinguishable and intermediate between the no-risk (L-P-) and both-stage (L+P+) risk environments (Fig 4a).

**Fig 4:**
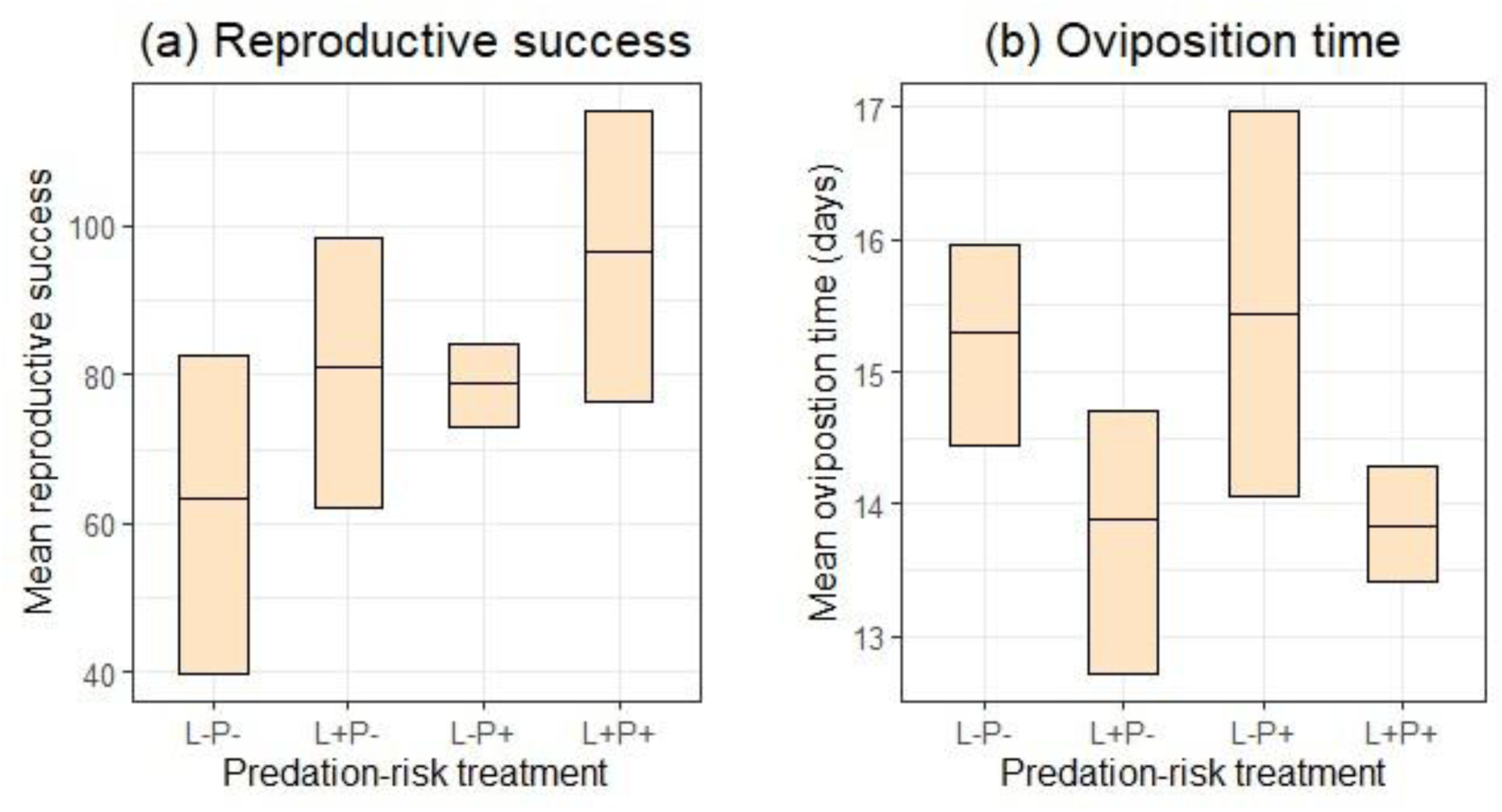
(a) Mean reproductive success and (b) average oviposition time data (mean + 95% bootstrap confidence intervals) for females across different risk treatments, i.e., no-risk (L-P-), risk during the larval stage (L+P-), the pupal stage (L-P+), and both the stages (L+P+). Data were collected from five cages per treatment over 24-26 days.

**Table 4:** Model estimates, their confidence intervals, and test statistics from linear mixed-effects models comparing mean reproductive success and average oviposition time of females from different risk treatments, i.e., risk vs. no-risk (contrast 1), risk during one stage vs. both the stages (contrast 2), risk during larval vs. pupal stage (contrast 3). Entries in bold denote statistically detectable effects.

| Trait | Estimate (orthogonal) contrasts | Estimate/ coefficient | 95% Conf. interval | t value | P value |
|---|---|---|---|---|---|
| Reproductive success | contrast 1<br>(L+P-, L-P+, L+P+) vs. L-P- | -5.527 | -11.379, 0.325 | -1.779 | 0.097 |
|  | contrast 2<br>(L+P-, L-P+) vs. L+P+ | 5.500 | -2.776, 13.776 | 1.252 | 0.231 |
|  | contrast 3<br>L+P- vs. L-P+ | -1.000 | -15.335, 13.335 | -0.131 | 0.897 |
| Oviposition time | contrast 1<br>(L+P-, L-P+, L+P+) vs. L-P- | 0.228 | -0.083, 0.540 | 1.378 | 0.189 |
|  | contrast 2<br>(L+P-, L-P+) vs. L+P+ | -0.273 | -0.714, 0.167 | -1.164 | 0.263 |
|  | contrast 3<br>L+P- vs. L-P+ | **0.776** | **0.012, 1.540** | **1.909** | **0.076** |

When pooled together, females from risk environments showed similar mean oviposition time to females from the no-risk environment (Table 4: Oviposition time, contrast 1). Also, within risk environments, females from single-stage and both-stage risk environments did not statistically differ in their oviposition time (Table 4: Oviposition time, contrast 2). However, average oviposition time detectably differed between the single-stage risk environments (Table 4: Oviposition time, contrast 3). Specifically, females exposed to risk exclusively during the larval stage laid eggs approximately 1-1.5 days earlier than females exposed to risk during the pupal stage (Fig 4b). Therefore, although no differences in oviposition time were detected among predation risk treatments (contrasts 1 & 2), means and 95% bootstrapped confidence intervals revealed that females from the both-stage risk environment—like those from the larval-risk environment—laid eggs 1-1.5 days earlier than those from the pupal-risk and no-risk environments (Fig 4b). Hence, exposure to risk during the larval stage, with or without the pupal stage, significantly advanced oviposition.

#### Energy reserves

On average, predation risk exposure during the immature stages increased energy content per unit mass in males but not in females (Table 5: Protein/Mass, Glycogen/Mass & Lipid/Mass, contrast 1). Early risk exposure increased protein reserves in males by 5-15 μg per unit mass (7-20%), glycogen reserves by 10-12 μg per unit mass (18-21%) and lipid reserves by 55-90 μg per unit mass (18-30%) (Fig 5). There were no stage-specific risk effects on glycogen and lipid reserves (Table 5: Glycogen/Mass & Lipid/Mass, contrasts 2 & 3). Compared with glycogen and lipid, protein reserves varied with the stage of risk exposure.

**Fig 5:**
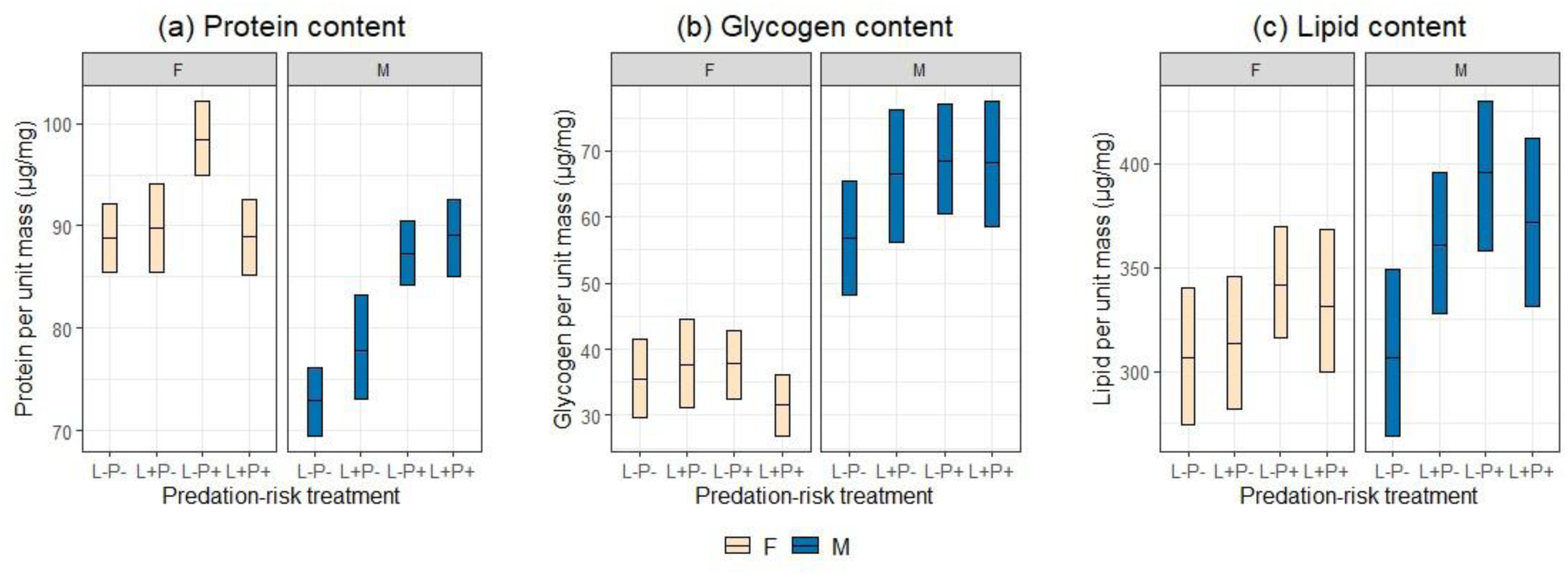
(a) Protein, (b) glycogen and (c) lipid content (mean + 95% bootstrap confidence intervals) of females (ivory) and males (blue) from different risk treatments, i.e., no-risk (L-P-), risk during the larval stage (L+P-), the pupal stage (L-P+), and both the stages (L+P+). Sample sizes: protein (F = 148, M = 149), glycogen (F = 146, M = 145) and lipid (F = 147, M = 148). Treatment-specific sample sizes are provided in Table 1.

**Table 5:** Model estimates, their confidence intervals, and test statistics from separate linear mixed-effects models for females and males, comparing the energy reserves across different risk treatments, i.e., risk vs. no-risk (contrast 1) risk during one stage vs. both the stages (contrast 2), risk during larval vs. pupal stage (contrast 3). Entries in bold denote statistically detectable effects.

| Trait | Estimate (orthogonal) contrasts | Female |  |  |  | Male |  |  |  |
|---|---|---|---|---|---|---|---|---|---|
|  |  | Estimate/ coefficient | 95% Conf. interval | t value | P value | Estimate/ coefficient | 95% Conf. interval | t value | P value |
| Protein / Mass | contrast 1<br>(L+P-, L-P+, L+P+) vs. L-P- | -1.044 | -2.222, 0.148 | -1.721 | 0.087 | **-3.103** | **-4.390, -1.805** | **-4.687** | **<0.0001** |
|  | contrast 2<br>(L+P-, L-P+) vs. L+P+ | -1.562 | -3.205, 0.065 | -1.866 | 0.064 | **2.103** | **0.301, 3.938** | **2.258** | **0.025** |
|  | contrast 3<br>L+P- vs. L-P+ | **4.393** | **1.432, 7.348** | **2.899** | **0.004** | **4.120** | **0.984, 7.362** | **2.526** | **0.012** |
| Glycogen / Mass | contrast 1<br>(L+P-, L-P+, L+P+) vs. L-P- | -0.294 | -1.561, 0.982 | -0.452 | 0.652 | **-2.516** | **-4.624, -0.417** | **-2.334** | **0.021** |
|  | contrast 2<br>(L+P-, L-P+) vs. L+P+ | -1.642 | -3.388, 0.090 | -1.843 | 0.067 | 0.647 | -2.230, 3.510 | 0.440 | 0.660 |
|  | contrast 3<br>L+P- vs. L-P+ | 1.133 | -2.058, 4.291 | 0.697 | 0.487 | 1.234 | -3.843, 6.297 | 0.475 | 0.635 |
| Lipid / Mass | contrast 1<br>(L+P-, L-P+, L+P+) vs. L-P- | -6.400 | -13.905, 1.132 | -1.661 | 0.099 | **-18.044** | **-26.500, -9.576** | **-4.162** | **<0.0001** |
|  | contrast 2<br>(L+P-, L-P+) vs. L+P+ | 1.153 | -9.298, 11.618 | 0.215 | 0.830 | -0.175 | -12.129, 11.726 | -0.029 | 0.977 |
|  | contrast 3<br>L+P- vs. L-P+ | 15.153 | -3.609, 33.869 | 1.578 | 0.117 | 19.347 | -1.641, 40.276 | 1.802 | 0.073 |

On average, males exposed to risk during both larval and pupal stages stored more protein per unit mass than those exposed during a single stage (Table 5: Protein/Mass, contrast 2). However, protein reserves varied largely between the single-stage risk environments. Males exposed to pupal-stage risk stored 9-10 μg (12%) more protein per unit mass than those exposed only during the larval stage (Fig 5a, Table 5: Protein/Mass, contrast 3), suggesting that the difference in protein reserves between the pooled single-stage group and both-stage group (contrast 2) was driven by the larval-risk group. Supporting this, means and bootstrapped confidence intervals showed that males exposed to risk during the pupal stage, with or without the larval stage, had 9-10 μg (12%) higher protein reserves than those exposed to larval-stage risk (Fig 5a).

Similarly, in females, pupal-stage risk increased protein reserves by 9 μg per unit mass, i.e., 9% more than in those exposed to larval-stage risk (Fig 5a, Table 5: Protein/Mass, contrast 3). Although the pooled effect of single-stage risk exposure did not significantly differ from that of both-stage exposure (Table 5: Protein/Mass, contrast 2), means and bootstrapped confidence intervals showed that protein reserves under pupal-stage risk were higher than under both-stage exposure. Overall, risk exposure during the pupal stage increased protein reserves by 9 μg per unit mass (9%) relative to all other treatments (Fig 5a).

In summary, males stored the most protein when exposed to risk during the pupal stage, alone or in combination with the larval stage, and maximised glycogen and lipid reserves regardless of the stage of risk exposure. In contrast, early-life risk did not affect overall energy reserves in females; only protein reserves were elevated in those from the pupal-risk environment.

### Effects of larval environment on traits till pupal stage

Approximately 98-99% of larvae survived in the risk environment until 95% of the population pupated, a rate comparable to that observed in the no-risk environment (Table 6: *z* = −0.978, *P* = 0.328). Although comparisons of means and their 95% bootstrapped confidence intervals suggested slightly reduced survival in the risk environment, the effect size was minimal (<1%) and biologically negligible (Fig 6a).

**Fig 6:**
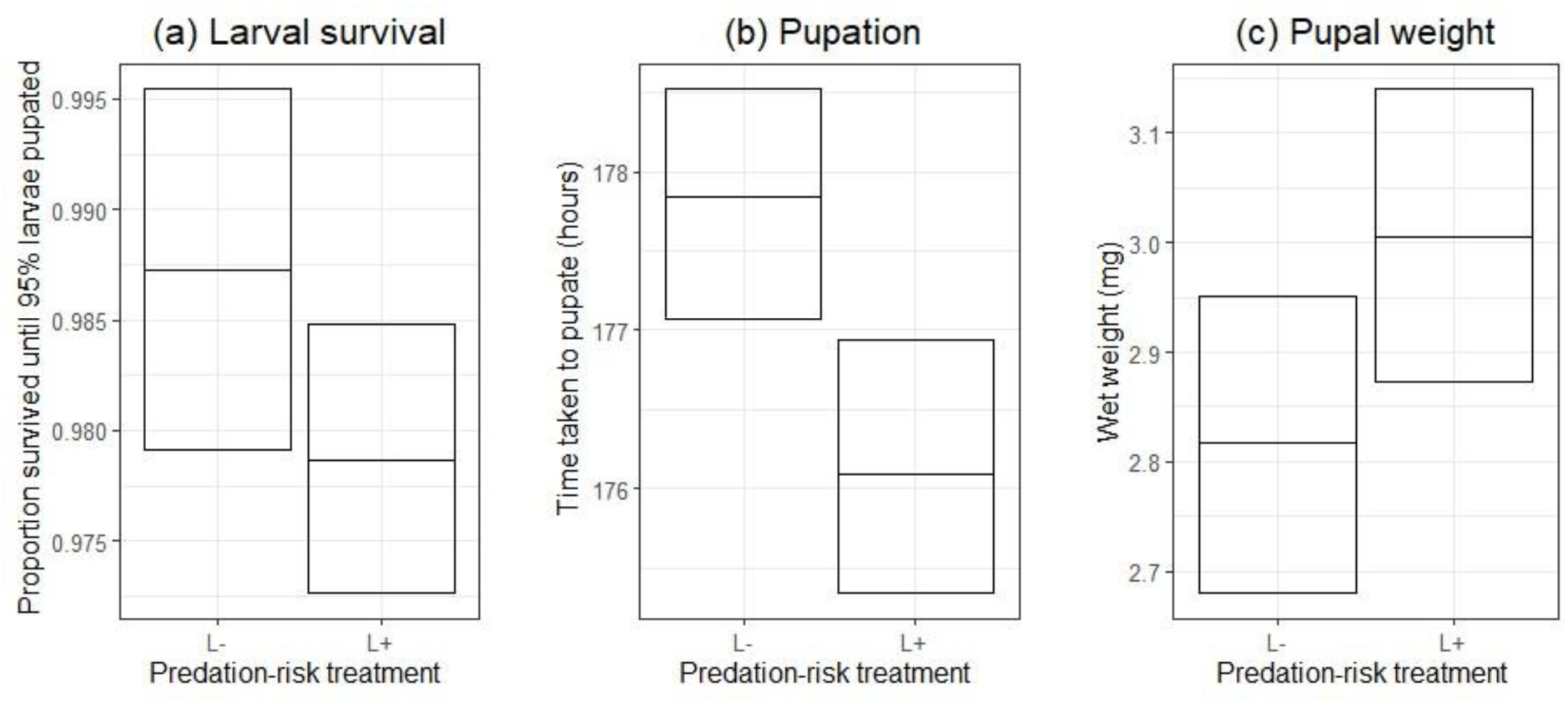
(a) Larval survival, (b) time to pupation, and (c) pupal weight data (mean + 95% bootstrap confidence intervals) from ‘no-risk’ (L-) and ‘risk’ (L+) larval environments. Sample sizes: survival (L- = 10 trays, L+ = 9 trays; 45 larvae/tray), pupation (L- = 432, L+ = 389) and weight (L- = 59, L+ = 60).

**Table 6:** Model estimates, their confidence intervals, and test statistics from a binomial mixed-effects model comparing the proportion of larvae surviving to the pupal stage, linear models comparing time to pupation and pupal mass, and linear mixed-effects models comparing energy reserves across different risk treatments, i.e., no-risk vs. risk during the larval stage. Entries in bold denote statistically detectable effects.

| Trait | Parameter | Estimate/ coefficient | 95% Conf. interval | t value/ z value | P value |
|---|---|---|---|---|---|
| Larval survival to pupation | Intercept (block1:no risk) | 4.304 | 3.590, 5.230 | 10.472 | <0.0001 |
|  | risk | -0.519 | -1.620, 0.508 | -0.978 | 0.328 |
| Time to pupation | Intercept (block1:no risk) | 176.423 | 174.362, 178.485 | 167.985 | <0.0001 |
|  | risk | **-1.816** | **-3.443, -0.189** | **-2.192** | **0.028** |
| Pupal weight | Intercept (block1:no risk) | 2.748 | 2.503, 2.992 | 22.261 | <0.0001 |
|  | risk | 0.188 | -0.091, 0.468 | 1.331 | 0.186 |
| Protein / Pupal mass | Intercept (block1:no risk) | 82.065 | 74.772, 88.993 | 20.525 | <0.0001 |
|  | risk | **-8.650** | **-16.772, -0.558** | **-2.090** | **0.040** |
| Glycogen / Pupal mass | Intercept (block1:no risk) | 43.586 | -2.467, 89.569 | 1.762 | 0.175 |
|  | risk | 1.035 | -6.800, 8.889 | 0.260 | 0.795 |
| Lipid / Pupal mass | Intercept (block1:no risk) | 548.190 | 189.829, 907.182 | 2.842 | 0.062 |
|  | risk | **-110.701** | **-193.922, -27.480** | **-2.623** | **0.010** |

Additionally, risk-exposed larvae pupated approximately two hours earlier than those in the no-risk environment (Fig 6b, Table 6: *t* = −2.192, *P* = 0.028), while no difference was detected in their pupal mass (2.8-3 mg) by the statistical model (Table 6: *t* = 1.331, *P* = 0.186). However, comparisons of means and confidence intervals suggested that risk-exposed individuals may have attained heavier pupal bodies than their no-risk counterparts (Fig 6c). This apparent discrepancy between the statistical model output and the inference drawn from bootstrapped confidence intervals highlights the need for further investigation.

Risk-exposed larvae also pupated with lower protein and lipid reserves per unit mass than larvae from the no-risk environment. Protein reserves were approximately 9 μg, i.e., 10% lower in pupae with larval risk exposure (Fig 7a, Table 6: *t* = −2.090, *P* = 0.040), while lipid reserves were roughly 110 μg, i.e., 20% lower than pupae from no-risk environment (Fig 7c, Table 6: *t* = −2.623, *P* = 0.010). In contrast, pupal glycogen reserves remained similar between risk and no-risk environments (Fig 7b, Table 6: *t* = 0.260, *P* = 0.795).

**Fig 7:**
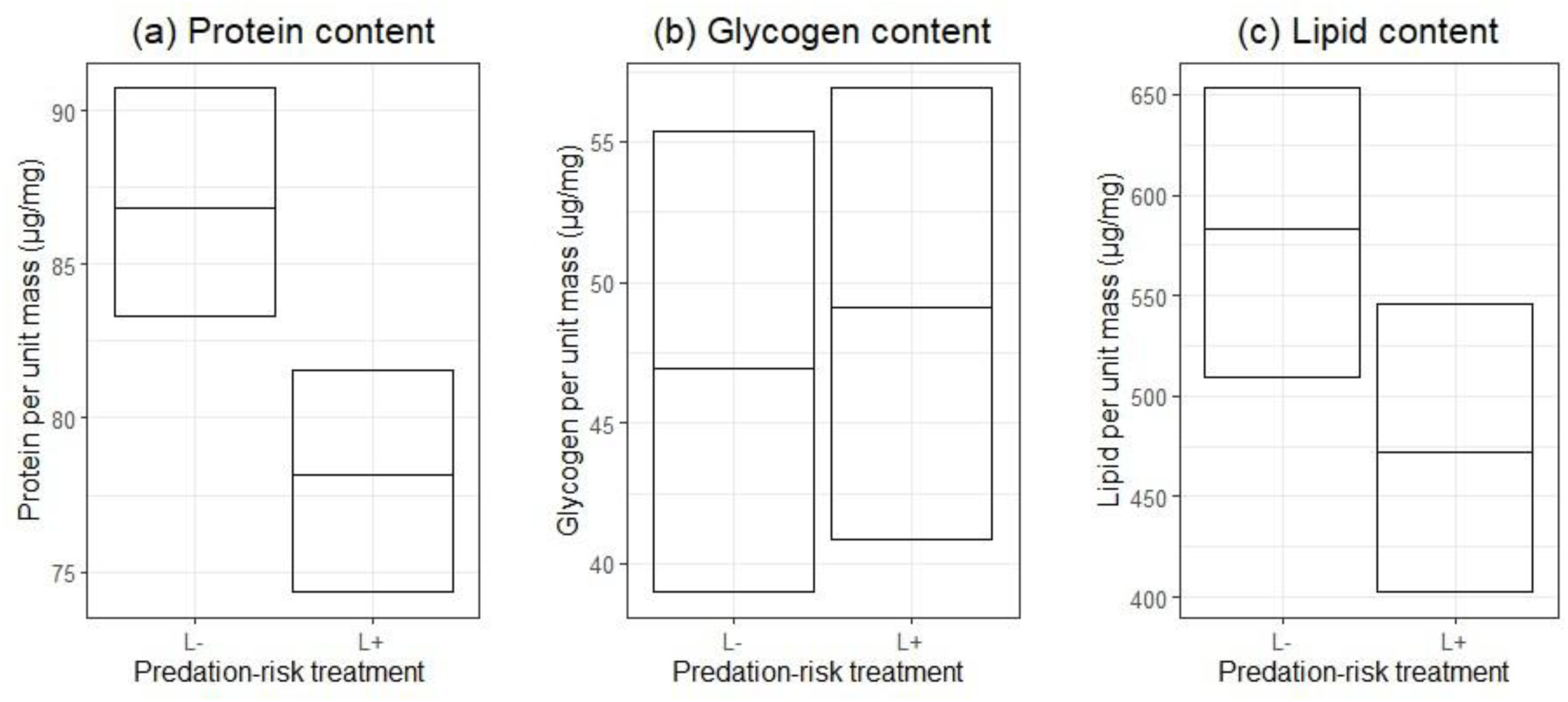
(a) Protein, (b) glycogen and (c) lipid content (mean + 95% bootstrap confidence intervals) of pupae from ‘no-risk’ (L-) and ‘risk’ (L+) larval environments. Sample sizes: protein (L- = 39, L+ = 40), glycogen (L- = 39, L+ = 40) and lipid (L- = 39, L+ = 39).

## Discussion

The multistage trade-off framework provided valuable insights into both cumulative and stage-specific effects of predation risk. Predation risk across multiple stages had varying impacts on different traits (Table 7a). Several developmental, survival-related, and reproductive traits reflected cumulative effects of risk exposure during the larval and pupal stages. In contrast, other traits showed similar responses regardless of whether the risk was encountered at a single stage or across multiple stages. These risk effects differed between males and females, a pattern observed in many taxa and often attributed to sex-specific behaviour and life-history strategies, with their physiological demands and other costs (Alto et al. 2005, Donelan and Trussell 2020, Gu et al. 2022, Maly 1970, Meuthen et al. 2019, Mirza et al. 2001, Lagos and Herberstein 2017, Shine et al. 2000, Wormington and Juliano 2014).

**Table 7:**
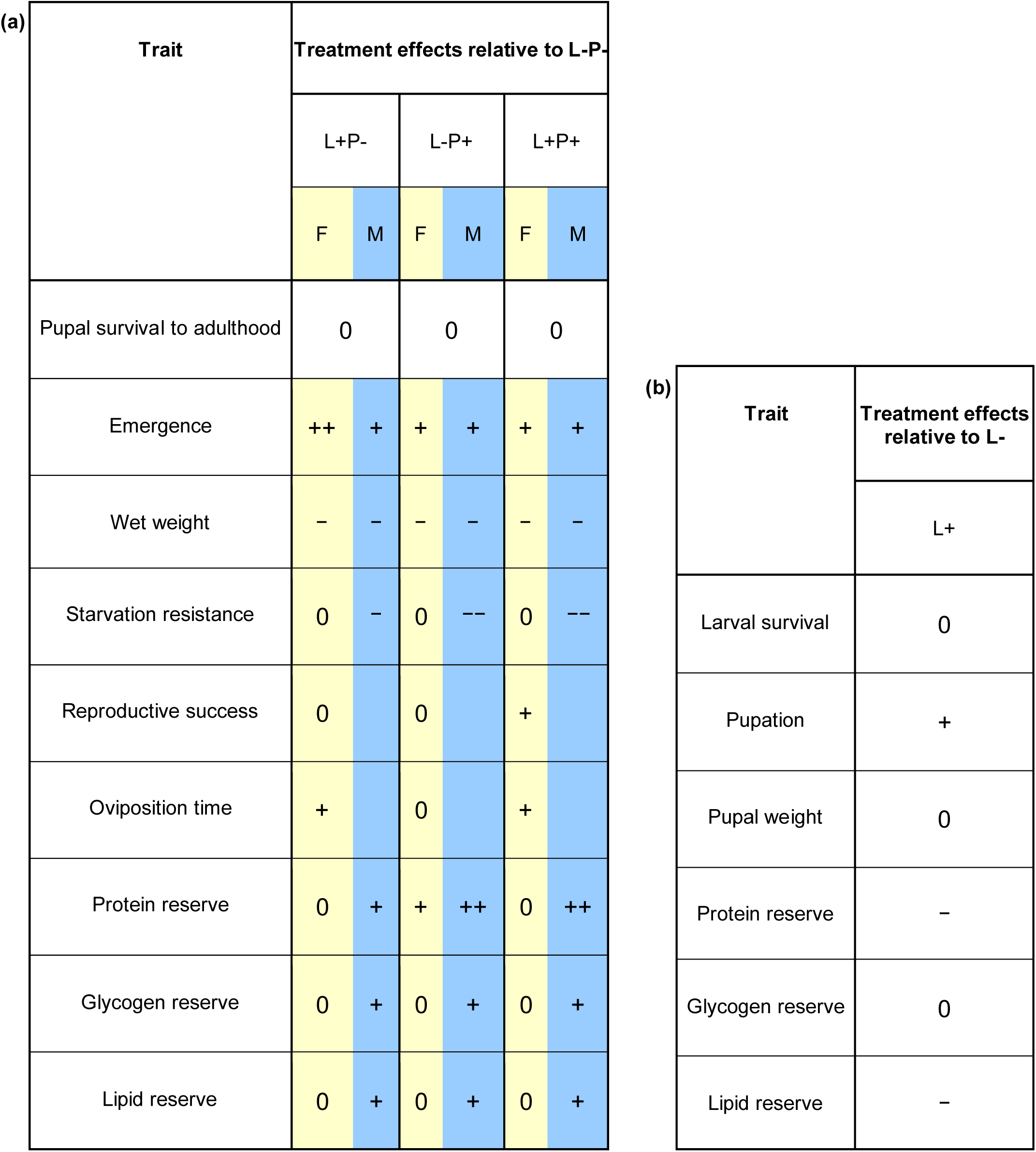
(a) Qualitative effects of larval and pupal predation risk relative to no-risk (L-P-), colour-coded by sex (ivory-yellow = females, blue = males). (b) Qualitative effects of larval-stage risk on traits up to the pupal stage relative to no-risk (L-). Symbols: 0 = no effect; +/− = positive/negative; ++/−− = stronger than +/−. See the ‘Statistical analyses’ section for inference details.

| (a) **Trait** | **Treatment effects relative to L-P-** |  |  |  |  |  |
|---|---|---|---|---|---|---|
|  | L+P- |  | L-P+ |  | L+P+ |  |
|  | F | M | F | M | F | M |
| Pupal survival to adulthood | 0 |  | 0 |  | 0 |  |
| Emergence | ++ | + | + | + | + | + |
| Wet weight | – | – | – | – | – | – |
| Starvation resistance | 0 | – | 0 | –– | 0 | –– |
| Reproductive success | 0 |  | 0 |  | + |  |
| Oviposition time | + |  | 0 |  | + |  |
| Protein reserve | 0 | + | + | ++ | 0 | ++ |
| Glycogen reserve | 0 | + | 0 | + | 0 | + |
| Lipid reserve | 0 | + | 0 | + | 0 | + |
| (b) **Trait** | **Treatment effects relative to L-** |  |  |  |  |  |
|  | L+ |  |  |  |  |  |
| Larval survival | 0 |  |  |  |  |  |
| Pupation | + |  |  |  |  |  |
| Pupal weight | 0 |  |  |  |  |  |
| Protein reserve | – |  |  |  |  |  |
| Glycogen reserve | 0 |  |  |  |  |  |
| Lipid reserve | – |  |  |  |  |  |

In males, survival under starvation was most reduced when both life stages were exposed to predation risk. Interestingly, males from the pupal-risk environment survived just as poorly, indicating that the pupal stage is particularly crucial in determining male survival. In contrast, female survival was not affected by risk exposure. Contrary to expectations, females exhibited enhanced reproductive success but only when both larval and pupal stages were exposed to risk. Additionally, oviposition occurred earlier when the larval stage experienced risk, regardless of pupal-stage exposure. Similarly, the larval environment appeared important for female emergence time. Females emerged fastest when exposed to risk during the larval stage alone, whereas pupal-stage exposure, alone or combined with larval-stage risk, led to a less pronounced acceleration in emergence.

Beyond these stage-specific outcomes, some traits responded similarly regardless of whether risk was experienced during one or both stages. In both sexes, body size (wet weight) was reduced in all risk environments. In males, emergence was accelerated, and energy reserves were elevated across all risk environments, although protein content peaked when the pupa was exposed to risk (L-P+ and L+P+). In females, however, there were no consistent changes in overall energy content except for increased protein levels in the L-P+ group. Importantly, predation risk had no detectable effect on pupal survival to adulthood in either sex.

Most traits did not align with the hypothesis predicting cumulative costs of predation risk across life stages. Only starvation resistance in males followed the expected pattern, and even then, males from the pupal-risk environment performed as poorly as those from the both-stage risk environment. This suggests that males manage survival costs more effectively when only the larval stage is exposed to risk. Likewise, the positive effects of predation risk, i.e., enhanced reproductive success and accelerated oviposition and emergence in females, were noticeably driven by the larval environment. As a short non-feeding stage, pupa likely offered limited opportunity to compensate for stress or invest in recovery, making it difficult to buffer the effects of risk exposure.

Overall, the results for body mass and male starvation resistance indicate costs of early predation risk. These findings are consistent with studies on various mosquito species, which have typically reported adverse effects of larval-stage predation risk (Alcalay et al. 2018, Beketov and Liess 2007, Bellamy and Alto 2018, Chandrasegaran and Juliano 2019, Ower and Juliano 2019, Roux et al. 2021, van Uitregt et al. 2012). At the same time, the positive effects observed in this study on development and energy reserves have also been documented, such as accelerated development in *Culex pipiens* and *Aedes triseriatus* (Alcalay et al. 2018, Ower and Juliano 2019) and elevated energy reserves in *Aedes aegypti* males exposed to early predation risk (Chandrasegaran et al. 2018). Similarly, the enhanced female reproductive success observed, though rare, parallels findings in other taxa. For example, female oriental fruit flies (*Bactrocera dorsalis*) that experience predation risk develop more rapidly with increased fecundity (X. Liu et al. 2024). These patterns across traits suggest trade-offs and broader life-history strategies, which are discussed further below.

The reduced body mass of fast-emerging males and females suggests a trade-off between life stages driven by antipredator responses of immature stages. Across taxa, animals vary in their developmental responses to predation risk (Benard 2004, Higginson and Ruxton 2010, Relyea 2007). In particular, individuals are expected to accelerate development during vulnerable stages to escape predation, though this may come at a cost of reduced body mass (Abrams and Rowe 1996, Arendt 1997). For example, *Daphnia pulex* neonates exposed to predator cues mature rapidly but develop into smaller adults (Dodson and Havel 1988). Similarly, exposure to cannibalistic conspecifics in invasive cane toads (*Rhinella marina*) induces hatchlings to speed up development, resulting in smaller tadpoles with lower survival rates (Crossland et al. 2024). In the present study, both males and females likely exhibited this trade-off, yet the consequences for their measured fitness components (starvation resistance and reproductive success) differed substantially.

Notably, male survival under starvation was reduced following early-life exposure to predation risk. In females, unlike in males, survival was unaffected by risk exposure. Moreover, the reduced body mass observed in females did not translate into immediate fitness costs. On the contrary, females from risk environments, despite having lower mass, exhibited better lifetime reproductive output than those from the no-risk environment. Females exposed to risk during both larval and pupal stages, in particular, laid more eggs and oviposited faster than females from the no-risk environment, a pattern that is rare but has also been observed in female oriental fruit flies that develop under predation risk and show increased fecundity (X. Liu et al. 2024). Together with the findings in males discussed below, these results suggest an investment trade-off shaped during development, prioritising reproduction over survival. However, offspring quality was not assessed and therefore warrants further study, as there may be costs in terms of offspring quality (Fleming and Gross 1990, Lack 1947, Smith and Fretwell 1974, Walsh and Reznick 2008).

While we could not measure male reproductive output, males from risk environments had elevated energy reserves. Specifically, protein content, which was elevated in all risk environments, was highest when the pupal stage was exposed to risk. High protein levels, for instance, have been associated with lekking behaviour in Mediterranean fruit fly males (Yuval et al. 1998). Glycogen and lipid reserves are linked to mate searching and mating success in male yellow dung flies, *Scathophaga stercoraria* (Blanckenhorn et al. 2003, Otronen 1995), while glycogen also fuels swarming activity in *Anopheles gambiae* (Maïga et al. 2012). Therefore, high energy content per unit mass in males from early-risk environments suggests they may possess greater reproductive potential despite reduced survival under starvation. Similarly, the high reproductive performance observed in females from both-stage risk environments may come at the cost of reduced survival in the wild, given their low body mass (Whitman 2008).

These patterns suggest a broader strategy beyond trade-offs between life stages: when exposed to early-risk environments, individuals may prioritise reproductive investment over survival, reflecting a reproduction-survival trade-off (Kirkwood and Rose 1991, Stearns 1989, Williams 1966). This trade-off is well-documented across insects, birds and mammals (Culina et al. 2019, Dobson and Jouventin 2010, Orell and Belda 2002, Partridge and Farquhar 1981). For example, in a population of red-billed choughs (*Pyrrhocorax pyrrhocorax*), females with enhanced reproductive output early in life tend to have shorter lifespans (Reid et al. 2003). Similarly, female rhesus macaques (*Macaca mulatta*) that invest heavily in offspring production during early adulthood exhibit reduced longevity (Blomquist 2009). These trade-offs are often observed under ecological conditions that reduce survival prospects, such as resource variability and predation (Gadgil and Bossert 1970, Javoiš and Tammaru 2004, Polak and Starmer 1998), a scenario particularly relevant for species with a fast pace of life or r-selected life histories (Hamel et al. 2010, Yuan et al. 2023). For instance, experimentally parasitised male *Drosophila nigrospiracula* with shorter lifespans show increased courtship effort compared with unparasitised males (Polak and Starmer 1998). The present study suggests that predation risk experienced early in life, not just during the adult stage, can initiate or modulate trade-offs between reproduction and survival. Such reproduction-survival trade-offs could be rigorously examined by tracking individual-level traits while accounting for among-individual variation in resource acquisition and allocation (van Noordwijk and de Jong 1986).

A unique aspect of this study was assessing the effects of risk exposure during the pupal stage. The reduced lifespan of starved males from the pupal-risk environment reveals that the pupal stage, though often overlooked, is a critical period during which risk exposure can lead to lasting physiological costs, likely because such costs cannot be offset in the absence of feeding. Furthermore, the effects of the pupal-risk environment―elevating protein content in both males and females, and increasing reproductive success in females when combined with larval risk exposure―indicate that the pupal stage may play a pivotal role in strategic allocation decisions (Boggs 2009). However, due to the ecology and life cycle of *Aedes aegypti*, it was challenging to isolate the pupal stage environment entirely. In the L-P+ environment, the last few hours (up to 24 hours for some) of the late larval stage were inadvertently exposed to the same conditions as the pupal stage. Despite this overlap, we believe that any effect in the L-P+ environment is driven mainly by the pupal stage. Nonetheless, future research utilising precise, stage-specific manipulations will be instrumental in disentangling the individual impacts of larval and pupal risk exposure.

We also examined the effects of the larval environment on pupal traits (Table 7b) to gain deeper insights into the role of pupa as the risk effects accumulate to influence the adult stage. Similar to accelerated adult emergence, pupation occurred earlier in larvae exposed to predation cues. This response is common in various insect species where larvae pupate early, with varying body mass, to evade a risky environment (Alcalay et al. 2018, Y. Liu et al. 2024, Xiong et al. 2015, Zhang et al. 2023; but see Hechtel and Juliano 1997, Martel et al. 2011, Russel and Cator 2022). These studies also report reduced survival to the pupal stage in the presence of predation cues. However, in this study, larval survival to pupation was unaffected by exposure to predation cues. Furthermore, despite the observed reduction in adult mass, we did not detect a corresponding decrease in pupal mass due to larval risk exposure. This suggests that the cost of rapid development manifests only during the pupal-to-adult transition: individuals that develop early under risk conditions are not lighter at pupation, but they emerge as lighter adults compared with those from no-risk environments.

As observed in adults, the effects of larval stress on pupal traits can also be sex-specific. For example, in the speckled wood butterfly (*Pararge aegeria*), increased larval rearing density reduces female pupal mass more than male pupal mass (Gibbs et al. 2004). Similarly, food stress decreases pupal mass only in female Glanville fritillary butterflies, *Melitaea cinxia* (Rosa and Saastamoinen 2017). Since we could not determine the sex of individuals at the pupal stage, effects observed in some pupal traits may have been driven primarily by one sex, while the absence of effects on other traits may reflect opposing responses in males and females. Nevertheless, the lack of an effect on pupal mass suggests that at least one sex, if not both, did not bear the cost of early pupation in the form of reduced body mass.

Additionally, pupal glycogen reserves remained unaffected, while lipid and protein reserves decreased due to larval risk exposure. This suggests that pupal energy reserves are compromised, at least in one sex, when larvae experience risk. However, pupae may conserve these reserves during metamorphosis, which could explain why adults from risk environments exhibited similar or higher energy content per unit mass compared with those from the no-risk environment. Alternatively, this increase in energy content per unit mass may reflect greater water loss during the pupal-to-adult transition, with risk-exposed individuals losing more water than those from no-risk environments, as the measurements were based on wet rather than dry weight.

Predation risk during early life stages can influence the complex energetics of metamorphosis. Although studies examining pupal energetics under ecologically relevant stressors are rare, these findings demonstrate the value of such research in uncovering compensatory mechanisms and energy allocation strategies that support stress resilience during the pupal stage.

### Conclusion

This study demonstrates that early-life stressors can impact multiple traits differently, with even the brief pupal stage playing a crucial role. Males suffered reduced lifespan under starvation, which was further reduced when the pupal stage was exposed to risk. The cumulative and stage-specific survival costs of predation risk, therefore, establish the pupal stage as a particularly vulnerable period in responding to early-life predation risk. In contrast, females showed increased reproductive success and shorter oviposition time, particularly when both larval and pupal stages were exposed to risk, without a reduction in lifespan under starvation.

Earlier emergence, accompanied by lighter body weight, in both males and females, suggests a potential trade-off between development and body size. In addition, the study suggests a broader life-history strategy shaped by early predation risk, characterised by a trade-off between reproduction and survival. The pupal stage likely plays a pivotal role in mediating allocation strategies within this trade-off, as risk exposure during pupation was associated with higher protein reserves (crucial for reproduction) in males and improved reproductive performance in females. These results underscore the importance of examining multiple life stages to reveal how even seemingly dormant stages shape life-history trade-offs and adult fitness.

## Acknowledgements

We acknowledge the Indian Institute of Science for its institutional and logistical support throughout this study. We thank Kanchana Gaonkar, Manvi Sharma, Karthikeyan Chandrasegaran, and Akshay P. for sharing their knowledge and expertise. We are grateful to our interns, Subhiksha, Gopika, Angela, and Mohit, for their dedicated assistance with the experiments and data verification. We also thank Dr Deepak Saini and his laboratory, along with Dr Dipshikha Chakravortty, for their technical guidance during the biochemical assays and for providing access to laboratory facilities. We extend our thanks to Ravi for his beautiful illustrations of the mosquito life stages.

This research received fellowship support from the Ministry of Education, Government of India, and grant support from the Department of Science and Technology–Science and Engineering Research Board (DST-SERB) under the Scientific and Useful Profound Research Advancement (SUPRA) programme (SPR/2020/000384). Additional financial support was provided through DST-FIST [Fund for Improvement of S&T Infrastructure; sanction number SR/FST/LSII-025/2009(C)], the DBT-IISc Partnership Program (Phase II; Department of Biotechnology, Ministry of Science and Technology, and Indian Institute of Science, India), CSIR (Council of Scientific and Industrial Research) 37(1636)/14/EMR-II), and the Indian Institute of Science.

The study received approval from the Institutional Animal Ethics Committee of the Indian Institute of Science (CAF/Ethics/835/2021) for mosquito breeding and experiments involving live insects.

